# Discovery of the apiosyltransferase, celery UGT94AX1 that catalyzes the biosynthesis of a flavone glycoside, apiin

**DOI:** 10.1101/2023.05.22.541790

**Authors:** Maho Yamashita, Tae Fujimori, Song An, Sho Iguchi, Yuto Takenaka, Hiroyuki Kajiura, Takuya Yoshizawa, Hiroyoshi Matsumura, Masaru Kobayashi, Eiichiro Ono, Takeshi Ishimizu

## Abstract

Apiose is a unique branched-chain pentose found in plant glycosides and a key component of the cell wall-polysaccharide pectin and other specialized metabolites. More than 1,200 plant-specialized metabolites contain apiose residues, represented by apiin, a distinctive flavone glycoside found in celery and parsley in the family Apiaceae. The physiological functions of apiin remain obscure, partly due to our lack of knowledge on apiosyltransferase during apiin biosynthesis. Here, we identified celery UGT94AX1 (AgApiT) as a novel apiosyltransferase, responsible for catalyzing the last sugar-modification step in apiin biosynthesis. AgApiT showed strict substrate specificity for the sugar donor, UDP-apiose, and moderate specificity for acceptor substrates, thereby producing various apiose-containing flavone glycosides in celery. Homology modeling of AgApiT with UDP-apiose, followed by site-directed mutagenesis experiments, identified unique Ile139, Phe140, and Leu356 residues in AgApiT, which are seemingly crucial for the recognition of UDP-apiose in the sugar donor pocket. Sequence comparison and molecular phylogenetic analysis of celery glycosyltransferases paralogous to AgApiT suggested that *AgApiT* is the sole apiosyltransferase-encoding gene in the celery genome. This is the first report on the identification of a plant apiosyltransferase gene that will enhance our understanding of the physio-ecological functions of apiose and apiose-containing compounds.

## Introduction

Apiose is a branched-chain aldopentose that is predominantly found in plants (Pičmanová and Møller, 2016), and rarely in lichens (Řezanka and Guschina, 2000), molds (Zheng et al., 1998) or bacteria (Smith and Bar-Peled, 2017). Apiose residues in plants are distributed in cell wall pectins and many specialized metabolites. Pectic rhamnogalacturonan II—widely present in streptophyte plants (O’Neill et al., 2004)—contains apiose residues in its side chains. Another apiose-containing pectin component, apiogalacturonan, is abundantly present in aquatic monocots, such as duckweeds, (Lemnoideae) (Avci et al., 2018) or seagrasses (Zosteraceae) (Gloaguen et al., 2010). The apiose content in the cell wall of these species reportedly reaches > 30% (Avci et al., 2018). Further, the level of apiogalacturonan correlates with plant growth capacity (Pagliuso et al., 2018). The diol structure of apiose residues in pectin is responsible for forming borate diester bonds and contributes to higher-order structures in the cell wall (Kobayashi et al., 1996; O’Neill et al., 1996).

Apiose residues are also found in specialized metabolites. Indeed, nearly 1,200 apiose-containing metabolites, including flavonoid and cyanogenic glycosides, have been reported (Pičmanová and Møller, 2016). Furthermore, they are distributed in at least 100 plant families, among which, certain plant species produce a high amount of these compounds, especially celery (*Apium graveolens*) and parsley (*Petroselinum crispum*) of the family Apiaceae. Flavonoid glycosides containing apiose residues accumulate in relatively high amounts in celery and parsley, accounting for 0.4‒1.5% and 1.5‒3.7 % of the dry weight, respectively (Lin et al., 2007; Boutshika et al., 2021). Among these flavonoid glycosides, apiin (apigenin-7-*O*-β-D-apiofuranosyl-(1→2)-β-D-glucopyranoside) is the most abundant. Luteolin-7-*O*-β-D-apiofuranosyl-(1→2)-β-D-glucopyranoside and chrysoeriol-7-*O*-β-D-apiofuranosyl-(1→2)-β-D-glucopyranoside are also biosynthesized in celery and parsley (Lin et al., 2007; Boutshika et al., 2021). In addition to Apiaceae, these compounds have also been observed in Asteraceae, Fabaceae, Plantaginaceae, and Solanaceae (Watson and Orenstein 1975; Kashiwagi et al., 2005). Further, apiose residues are sporadically found in many plant species, but their relatively limited distribution suggests convergent evolution. Thus, apiose or apiose-containing compounds have seemingly evolved independently in different plant species and possess lineage-specific physiological roles, presumably as evolutionary consequences of local adaptation (Pichersky and Raguso, 2018).

Apiin is one of the first flavonoid glycosides initially isolated in 1843 from parsley and celery (Braconnot, 1843). These plants have been used for traditional insecticides and fungicides. In 1901, apiin was shown to contain apiose residues (Vongerichten, 1901). The apiose residue was bound to the glucose moiety of apigenin 7-*O*-glucoside. Recently, much attention has been paid to the health benefits of apiin/apigenin owing to their anti-inflammatory, antidepressant (Salehi et al., 2019), and antioxidant activities (Li et al., 2014; Epifanio et al., 2020). Moreover, apiin reportedly contributes to various stress responses in plants, i.e., 1) UV irradiation of cultured parsley cells induces the accumulation of apiin (Gardiner et al., 1980: Eckey-Kaltenbach et al., 1993), and 2) it is found in winter-hardy plants, including celery and parsley (Watson and Orenstein, 1975). Interestingly, while some insects use Apiaceae, Rutaceae, or Solanaceae plants for larval feeding and oviposition (Ehrlich and Raven, 1967), certain species expressed apiose-containing compounds show moderate deterrent activity for female oviposition by the swallowtail butterfly (*Papillo xuthus*) (Ono et al., 2004) or the serpentine leafminer fly (*Liriomyza trifolii*) (Kashiwagi et al., 2005), suggesting that apiin has an ecological role in plant-insect interactions. However, the relationship between apiin structure and function remains unclear. It is also unresolved how the terminal apiose portion contributes to the function of apiin. To address these unsolved questions, it is necessary to identify the genes encoding enzymes involved in apiin biosynthesis and to characterize their biochemical properties. Once these are identified, functional analysis of their defective mutant in plants will help to clarify the functions of apiin.

Most enzymes involved in the biosynthesis of apiin (Supplemental Figure S1) have been identified in celery. Thus, the chalcone synthase (*CHS*), chalcone isomerase (*CHI*), and flavone synthase I (*FNSI*) genes involved in the biosynthesis of apigenin—the aglycone portion of apiin—have been identified (Yan et al., 2014). However, the celery glucosyltransferase (*GlcT*) and apiosyltransferase (*ApiT*) genes that catalyze the formation of the disaccharide (apiosyl β1,2-glucoside) at the 7-*O* position of apigenin, have not yet been identified. The enzymes GlcT and ApiT, which are responsible for apiin synthesis, are expectedly UDP-sugar dependent glycosyltransferases (UGTs) of the GT1 family, which is an enzyme superfamily in plant genomes. Specifically, UGT is known to transfer sugar from a UDP-sugar (donor substrates) to various flavonoids and their glycosides (acceptor substrates) (Yonekura-Sakakibara and Hanada, 2011). Several GlcTs, which transfer glucose to apigenin 7-*O*-β-glucoside have been identified in other plant species but not in celery (Noguchi et al., 2009; Su et al., 2018).

In contrast, no *ApiT* gene involved in the biosynthesis of any apiose-containing compounds, including apiin, has been identified in any plant species, although ApiT activity (UDP-Api: flavone apiosyltransferase, EC 2.4.2.25) has been detected in parsley (Ortmann et al., 1970; Ortmann et al., 1972). Therefore, the ultimate goal of this study was to identify the apiosyltransferase gene involved in apiin biosynthesis. It will contribute to clarifying the structure and function relationship of apiin and the reason why celery and parsley produce large amounts of apiin. However, there are at least two difficulties in identifying ApiT genes. First, comparative genetic/genomic information is scarce, as apiin is a specialized flavone glycoside produced by few plant species but not by well-known model plants, such as Arabidopsis or rice. Second, the sugar donor substrate, UDP-Api, is not commercially available due to poor product stability. Recently, we successfully developed a method for stabilizing UDP-Api using bulky cations, such as triethylamine, as counter ions (Fujimori et al., 2019). This enabled us to produce UDP-Api for evaluation of ApiT activity. Then, using UDP-Api prepared by this method, we detected apiin-synthesis ApiT activity in crude parsley extracts (Fujimori et al., 2019). In addition, the celery genome has been sequenced recently (Li et al., 2020). These research breakthroughs prompted us to screen the apiin-synthesis ApiT gene in celery.

## Results

### Screening for candidate AgApiT-encoding genes

Celery apigenin 7-*O*-glucoside: apiosyltransferase (AgApiT) is an inverting glycosyltransferase because the linkages of the apiosyl anomers in UDP-Api and apiin are α and β, respectively. Further, AgApiT is supposed to belong to the glycoside-specific glycosyltransferases (GGTs) within UGTs, because the apiose residue is bound to the glucose residue of apigenin 7-*O*-glucoside by a β1-2 linkage. GGTs catalyze the transfer of additional sugar residues to the sugar moiety of flavonoid glycosides, resulting in disaccharides with a β1-2 or a β1-6 linkage (Frydman et al., 2004; Ono et al., 2020). Further, GGTs are mainly composed of UGT79, UGT91, and UGT94 in orthologous group 8 (OG8) (Yonekura-Sakakibara and Hanada, 2011). In celery, apiin is predominantly produced during early leaf development (Yan et al., 2014). Therefore, we attempted to identify the gene encoding AgApiT within celery genome and RNA-seq data, based on the following two criteria for *AgApiT* screening: 1) structural similarity to GGT and 2) co-expression with other apiin biosynthesis-related enzymes during early leaf development.

We found at least 26 GGT (UGT79, 91, and 94) genes in the celery genome data (Li et al., 2020) (Figure 1). Further, among these GGT genes, 10 were expressed in public RNA-Seq data (SRR1023730) (Table 1). The sequence of Agr35256, found in the genomic database (Li et al., 2020), matched with two distinct transcript sequences predicted from the RNA-Seq data. The two transcripts were designated as Agr35256-1 and Agr35256-2 (Table 1). Agr35256-1, which has a high read count in the RNA-seq data of celery leaves, was selected as one of the most likely gene candidates encoding AgApiT (Table 1) because the production of apiin is very high in celery leaves (Lin et al., 2007). Because the conditions of the RNA-Seq data SRR1023730 were not described in the database, we could not conclude Agr35256-1 was a good candidate for the apiin-synthetic apiosyltransferase gene. Therefore, we extracted RNA from a 2.0 cm long true leaf, in which apiin is synthesized in relatively high amounts (Figure 2A), and RNA-Seq analysis was performed thrice times (DRR396538, DRR396539, and DRR396540). The TPM values in each RNA-Seq analysis for the 10 GGT genes as shown in Table 1 indicated that the TPM value of Agr35256-1 was by far the highest compared to the others. Therefore, Agr35256-1 was identified as a strong candidate gene for apiin-biosynthetic apiosyltransferase.

**Figure 1.**
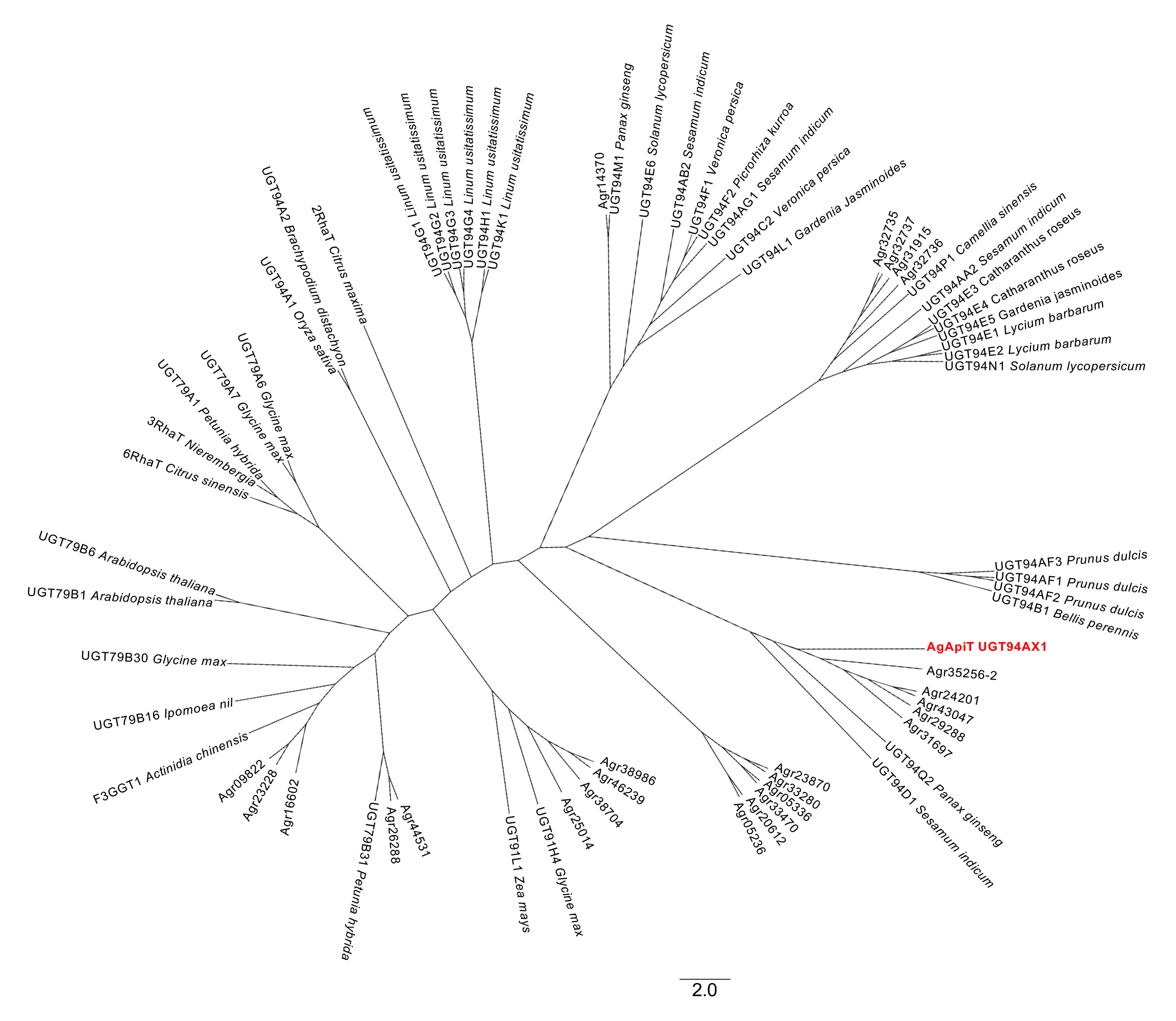
Phylogenetic tree of GGTs including AgApiT. The sequences were obtained from UGT Nomenclature Committee Website and celery DB (Li et al., 2020). Twenty-six celery GGTs are shown in AGR. AgApiT was shown in red.

**Figure 2.**
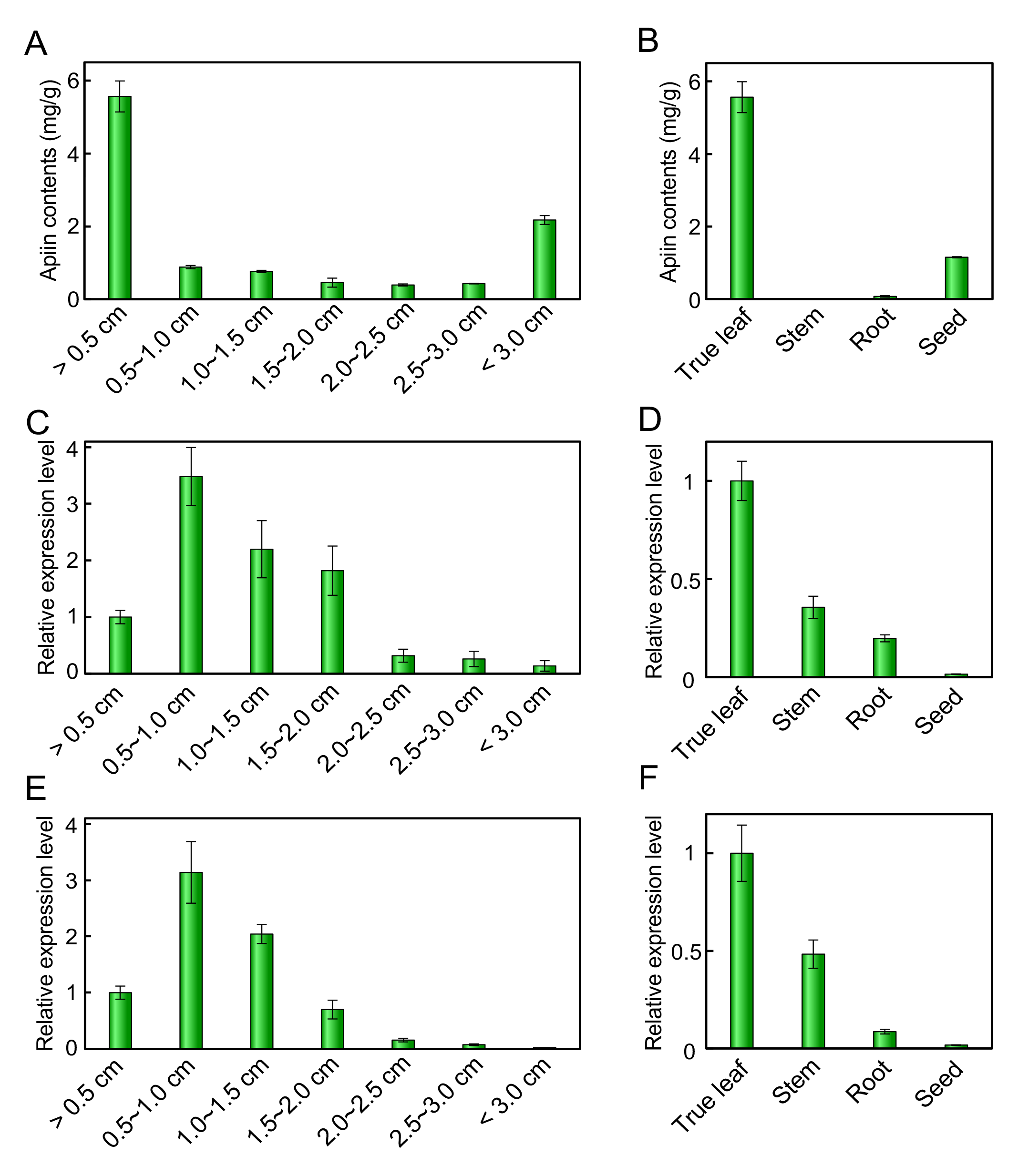
Apiin contents and expression of *AgApiT (UGT94AX1)* in leaf developmental stages and some tissues. Apiin contents (A, B) and quantitative RT-PCR analysis of *AgApiT (UGT94AX1)* (C, D) and *AgFNSI* (E, F) of celery true leave developmental stages (A, C, and E) and some tissues (B, D, and F). Apiin contents (mg) is shown per gram of each celery sample. The values on horizontal axis in (A, C, and E) indicate the longitudinal length of true leave. Each bar represents mean values and standard deviations from three biological replicates.

**Table 1.** Celery GGTs in OG8 and their expression level estimated using an RNA-Seq database.

| Gene ID | UGT family | TPM |  |  |  |
| --- | --- | --- | --- | --- | --- |
|  |  | SRR1023730 | DRR396538 | DRR396539 | DRR396540 |
| Agr35256-1 | UGT94 | 37.18 | 208.28 | 132.82 | 214.80 |
| Agr05236 | UGT94 | 14.34 | 0.04 | 0.29 | 0.14 |
| Agr14370 | UGT94 | 11.07 | 2.75 | 2.59 | 3.26 |
| Agr35256-2 | UGT94 | 9.41 | 3.74 | 1.88 | 3.77 |
| Agr23228 | UGT79 | 7.85 | 17.87 | 24.36 | 20.15 |
| Agr38074 | UGT91 | 5.86 | 0.25 | 0.21 | 0.19 |
| Agr09822 | UGT79 | 5.66 | 2.55 | 2.28 | 3.06 |
| Agr16602 | UGT79 | 4.68 | 6.41 | 2.82 | 5.55 |
| Agr46239 | UGT91 | 3.65 | 0.87 | 1.43 | 1.16 |
| Agr31697 | UGT94 | 2.03 | 4.40 | 3.92 | 3.14 |

The amino acid sequences of celery GGTs in OG8 were collected using the RNA-Seq database (SRR1023730). Their gene IDs, UGT families, and TPM scores are shown. Gene IDs have been provided according to the CeleryDB (Li et al., 2020).

The Agr35256-1 gene, which contains a complete open reading frame of UGT with the C-terminal PSPG (plant secondary product glycosyltransferase) motif for UDP-sugar donor recognition (Offen et al., 2006, Ono et al., 2010; Supplemental Figure S2), was assigned as UGT94AX1 by the UGT nomenclature committee (Mackenzie et al., 1997), and had an estimated molecular mass and an isoelectric point of 49 kDa and 5.7, respectively. These values are consistent with the previously reported molecular mass of 50 kDa and an isoelectric point of 4.8 for partially purified ApiT from parsley (Ortmann et al., 1972). In the GGT phylogenetic tree constructed, celery UGT94AX1 co-clustered with five celery paralogous GGT genes and was close to PgUGT94Q2 from Panax ginseng (Jung et al., 2014) and to SiUGT94D1 from sesame (Noguchi et al., 2008), both of which also forms β1-2 glucoside linkages (Figure 1). Thus, UGT94AX1 appeared to be the most likely candidate for AgApiT, which forms a β1-2 apiosyl linkage in apiin.

### *UGT94AX1* expression profile during celery development

In celery, apigenin—the aglycone of apiin—is synthesized at an early stage of leaf development (Yan et al., 2014). True leaves of the celery were used in this study (cultivar: New Cornell 619) and divided by different developmental stages, namely, 1: less than 0.5 cm, 2: 0.5 to 1 cm, 3: 1 to 1.5 cm, 4: 1.5 to 2 cm, 5: 2 to 2.5 cm, 6: 2.5 to 3 cm, and 7: more than 3 cm; apiin content was high in stage 1 and decreased thereafter (Figure 2A). Among the true leaves, stems, roots, and seeds, apiin content was highest in the true leaves (Figure 2B). Here, we investigated the expression profile of *UGT94AX1* and *AgFNSI*, which are the known apiin biosynthesis-related enzyme genes (Supplemental Figure S1), in true leaf developmental stages and some tissues. The expression of *UGT94AX1* increased in the early leaf-developmental stages, peaked at developmental stage 2, and decreased from stages 3 to 7 (Figure 2C). Among some tissues, *UGT94AX1* was highly expressed in true leave and less in other tissues (Figure 2D). A similar expression pattern was recorded for *AgFNSI* (Figures 2E and 2F). The apiin content and the expression level of *UGT94AX1* roughly coincide, suggesting that *UGT94AX1* was involved in the biosynthesis of apiin. Our findings thus showed that *UGT94AX1* was a candidate gene for apiin biosynthesis-related apiosyltransferase. Note that apiin contents were high in the seeds and in the main leaves less than 0.5 cm despite the low expression of the enzyme genes. It can be assumed that apiin was already biosynthesized in these two samples by the time they were collected.

### Evaluation of apigenin 7-*O*-glucoside: ApiT activity of UGT94AX1

Gene expression profiling was followed by an assessment of apigenin 7-*O*-glucoside: ApiT activity of UGT94AX1 (Figure 3A). The recombinant protein fused with the ProS2 tag was expressed in *Escherichia coli*. The purified protein was digested with a protease to remove the ProS2 tag eluted as a 49 kDa protein, which corresponded to the calculated molecular mass (Figure 3B). When this recombinant protein was incubated with 50 μM apigenin 7-*O*-glucoside in the presence of 0.5 mM UDP-Api, a new product was observed, with a retention time of 14.0 min on reversed-phase chromatography (Figure 3C). The levels of this enzyme product increased in an enzyme dose-and incubation time-dependent manner. The elution time of the product was identical to that of the authentic apiin. Therefore, we concluded that these results clearly showed that the UGT94AX1 protein has apigenin 7-*O*-glucoside: ApiT activity resulting in the production of apiin *in vitro*. Hereafter, UGT94AX1 is referred to as AgApiT. This is the first plant ApiT to be identified.

**Figure 3.**
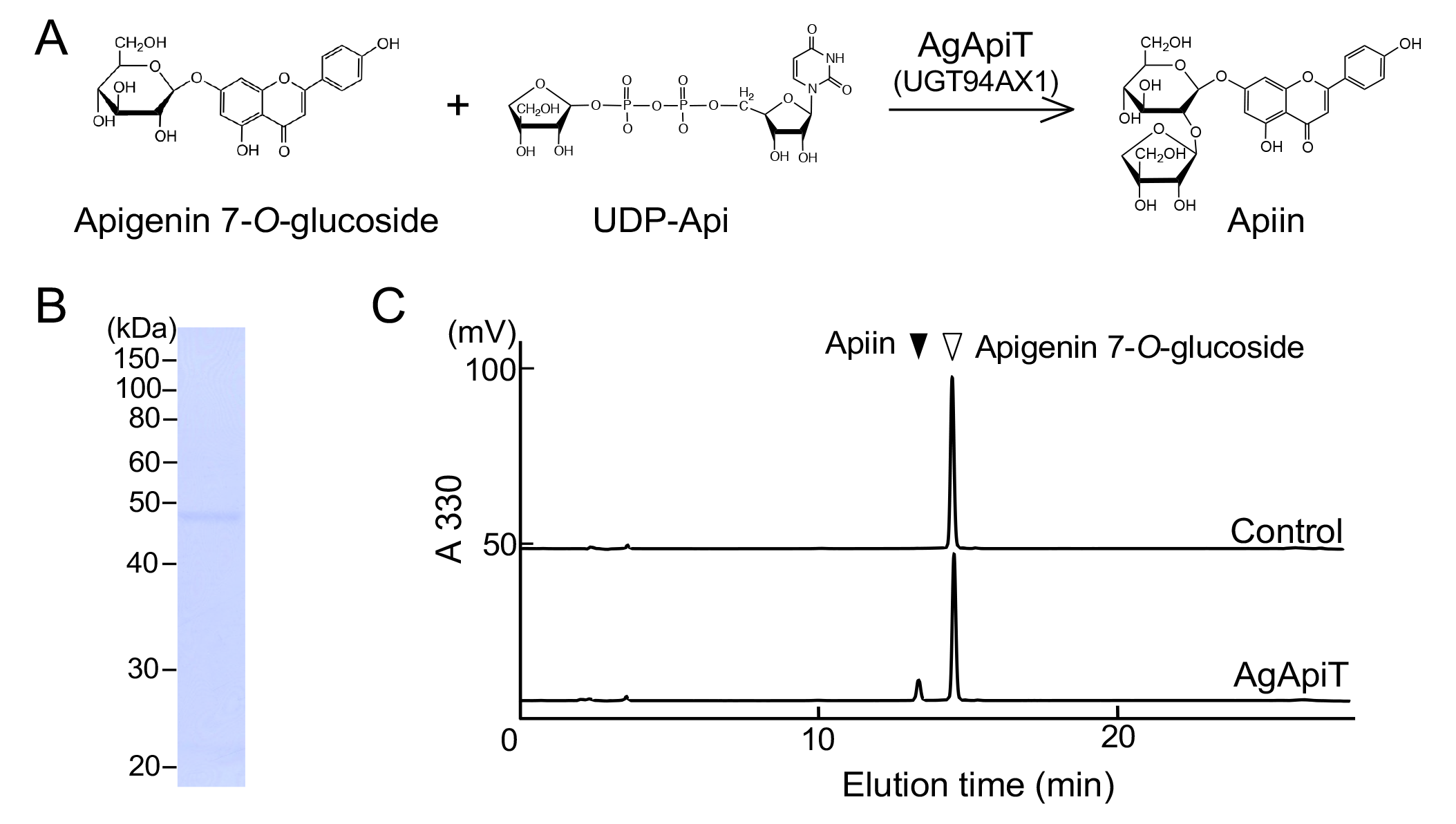
Apigenin 7-*O*-glucoside: apiosyltransferase activity of AgApiT. (A) The reaction of apigenin 7-*O*-glucoside: apiosyltransferase. (B) SDS-PAGE of the purified recombinant AgApiT. (C) Apiin-synthetic apiosyltransferase activity of the purified AgApiT. The substrate, apigenin 7-*O*-glucoside, yielded a single peak (upper panel). The apiosyltransferase activity of AgApiT resulted in the production of apiin as a product when it was incubated with 50 μM apigenin 7-*O*-glucoside and 0.5 mM UDP- Api at 23°C for 2 h (lower panel).

### Substrate specificity of AgApiT for sugar donors and acceptors

The sugar donor specificity of AgApiT was investigated using each of the 10 sugar nucleotides (UDP-sugars) with apigenin 7-*O*-glucoside as a sugar acceptor. The enzymatic reaction product was generated only when AgApiT was reacted with UDP-Api. Thus, the donor specificity of AgApiT was strictly specific to UDP-Api (Figure 4A). The *K*_m_ and *k*_cat_ values of AgApiT for UDP-Api were 8.6 ± 0.6 μM and (0.65 ± 0.01) × 10^-3^ s^−1^, respectively (Figure 4C). These values were comparable to those of previously characterized UGT94 enzymes specialized for different metabolites with branched sugar moieties in tea and sesame (Ohgami et al., 2015; Ono et al., 2020).

**Figure 4.**
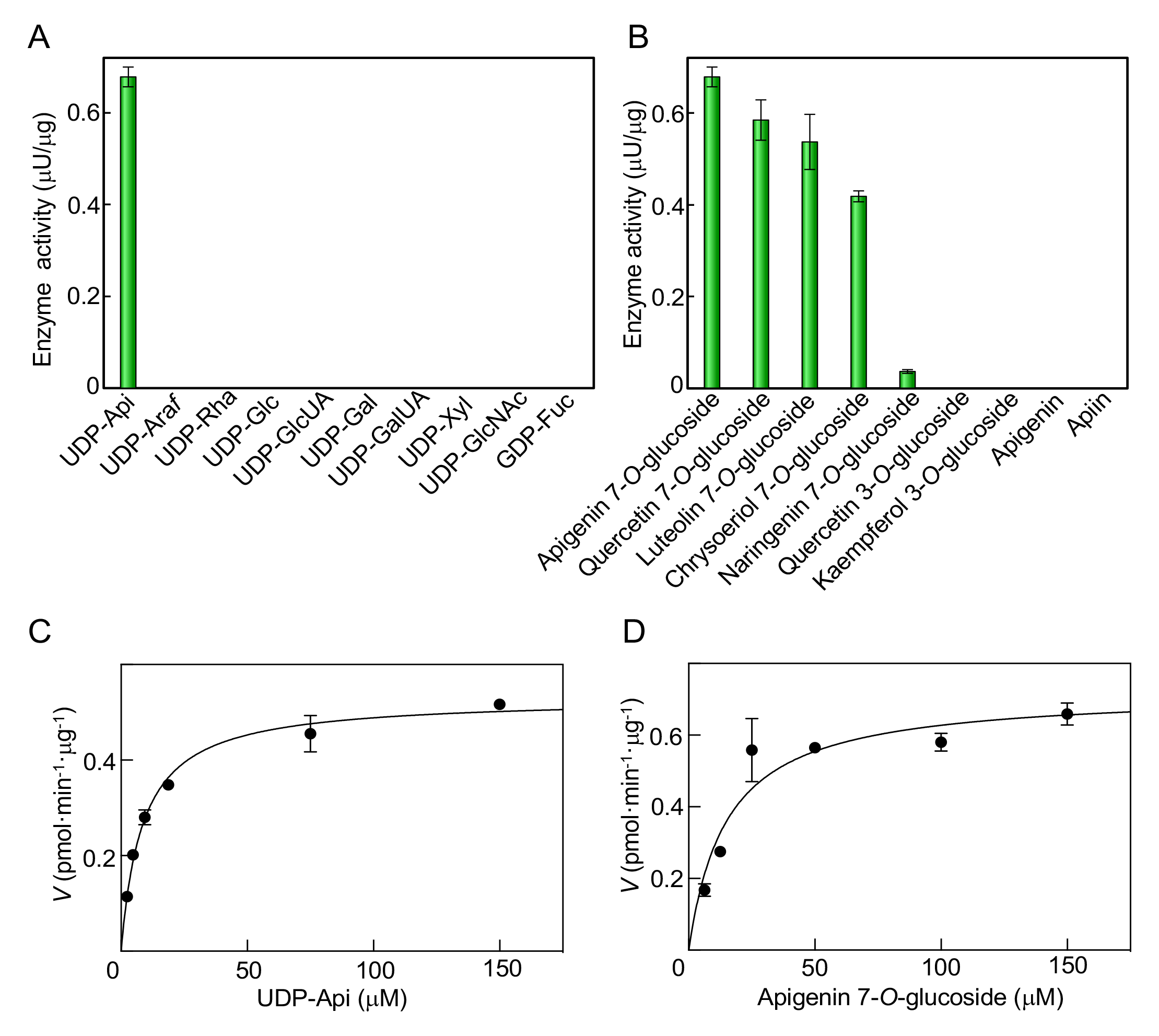
Substrate specificity and kinetic parameters of AgApiT. (A) Donor substrate specificity of AgApiT. The purified recombinant AgApiT was reacted with 50 μM apigenin 7-*O*-glucoside and 1 mM of each sugar nucleotide. (B) Acceptor substrate specificity of AgApiT. The purified recombinant AgApiT was reacted with 50 μM of each acceptor substrate and 1 mM UDP-Api. The activities are presented as mean values with standard errors of three independent samples. Michaelis-Menten plots of AgApiT for (C) UDP-Api and (D) apigenin 7-*O*-glucoside. The mean values with standard deviation (of three independent samples) of the velocity at each substrate concentration were plotted. The *K*_m_ and *k*_cat_ values were calculated from the nonlinear regression curve shown using the solid line.

Subsequently, the acceptor specificity of AgApiT was investigated using several flavonoid compounds. AgApiT reacted with 7-*O*-glucosides of flavone or flavonol (apigenin 7-*O*-glucoside, quercetin 7-*O*-glucoside, luteolin 7-*O*-glucoside, and chrysoeriol 7-*O*-glucoside) but negligibly with flavanone 7-*O*-glucoside (naringenin 7-*O*-glucoside) (Figure 4B). The double bond between C-2 and C-3 of flavone or flavonol, which is missing in flavanone, seems critical for substrate recognition of this enzyme. In contrast, AgApiT failed to react with flavonoide 3-*O*-glucosides, apiin, or its aglycone, apigenin (Figure 4B). We conclude that AgApiT preferentially reacts with 7-*O*-glucosides of flavone or flavonol. The *K*_m_ and *k*_cat_ values of AgApiT for apigenin 7-*O*-glucoside were 15 ± 3 μM and (0.88 ± 0.05) × 10^-3^ s^−1^ (Figure 4D). These values for the sugar acceptor are comparable to those obtained for previously characterized UGT94 enzymes (Ohgami et al., 2015; Ono et al., 2020). Altogether, these results support the idea that AgApiT is a UDP-Api-specific glycosyltransferase for flavone 7-*O*-glucosides, presumably producing apiin.

### Sequence comparison among GGTs and AgApiT

AgApiT shows strict sugar donor specificity, reacting only with UDP-Api (Figure 4A). To identify the amino acid residues responsible for the unique specificity of AgApiT, the structural features were investigated by comparing amino acid sequences among biochemically characterized GGTs (Supplemental Figure S2). Several biochemically characterized GGTs and their substrates (sugar donor and acceptor), and the amino acid sequences of sites for recognition of sugar donors, are summarized in Table 2.

**Table 2.**
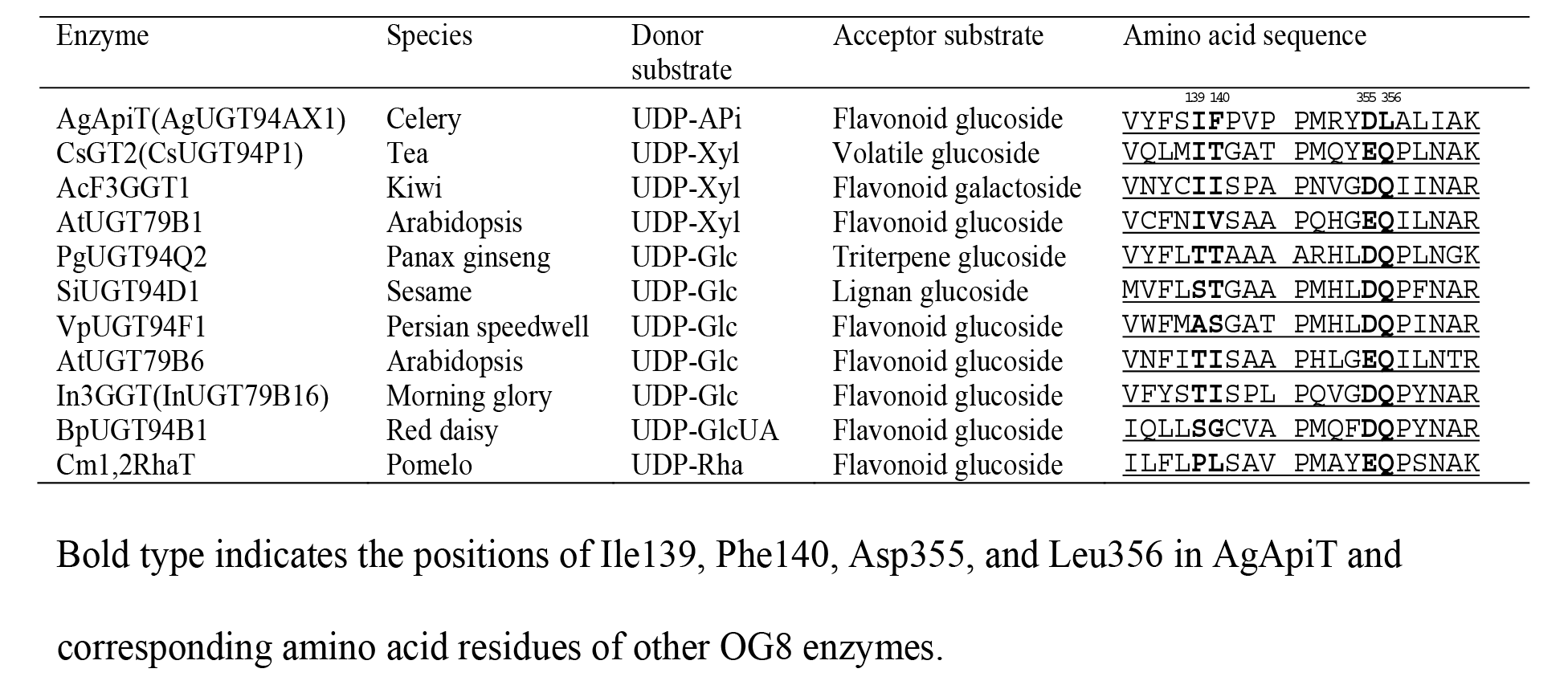
Substrate specificity and amino acid sequence comparison among GGTs in OG8.

A previous study showed that substituting Ile142 of tea CsGT2/UGT94P1 xylosyltransferase (XylT) with a Ser residue altered the donor specificity so that CsGT2-I142S utilized UDP-Glc instead of UDP-Xyl (Ohgami et al., 2015). Among the GGTs present in OG8, this Ile142 residue is uniquely conserved only in XylTs (Table 2) and is not found in other GlcTs (Frydman et al., 2004; Morita et al., 2005; Sawada et al., 2005; Jung et al., 2014). Thus, this conserved Ile residue among the XylTs was required to recognize the xylose moiety of UDP-Xyl in CsGT2. Furthermore, this Ile residue was also found in AgApiT (Ile139). According to the model structure of CsGT2, Thr143 adjacent to Ile142 is also part of the sugar recognition pocket (Ohgami et al., 2015). The corresponding Phe140 of AgApiT is a unique amino acid at this position. Therefore, Ile139 and Phe140 are candidate amino acid residues involved in apiose specificity in AgApiT.

In the crystal structure of flavonol 3-*O*-GlcT (VvGT1/UGT78A5) isolated from grape, the side chains of Asp374 and Gln375 independently form hydrogen bonds with 3- and 4-OH, and 3-OH of the glucose moiety in UDP-Glc (Offen et al., 2006). Moreover, structural modeling of CsGT2 also suggested that Glu369 and Gln370 bind to the xylose residue of the specific sugar donor, UDP-Xyl, via hydrogen bonding in the same manner (Ohgami et al., 2015). The corresponding amino acid residues in AgApiT are Asp355 and Leu356 (Table 2). Among biochemically characterized GGTs (Table 2) and the 26 celery GGTs (Figure 1), only AgApiT has Leu356, while others have a conserved Gln residue in this position. This unique amino acid polymorphism in AgApiT indicates that in addition to Ile139 and Phe140, Leu356 is another candidate residue underlying the specificity for the recognition of UDP-Api.

### Amino acid residues involved in the recognition of UDP-Api by AgApiT

To assess the enzymatic impact of Ile139, Phe140, or Leu356 of AgApiT on the specificity for UDP-Api, we evaluated the enzymatic activity of AgApiT mutants produced by site-directed mutagenesis. I139V showed 30% of the activity of wild-type AgApiT, whereas I139T and I139S showed no trace of enzyme activity (Figure 5A). Meanwhile, F140T, F140I, or F140V did not show any ApiT activity (Figure 5A). In turn, L356Q showed 15% of the activity of the wild-type enzyme, while neither L356I nor the L356T showed any enzyme activity (Figure 5A). Altogether, these data indicated that Ile139, Phe140, and Leu356 of AgApiT are required for complete ApiT activity.

**Figure 5.**
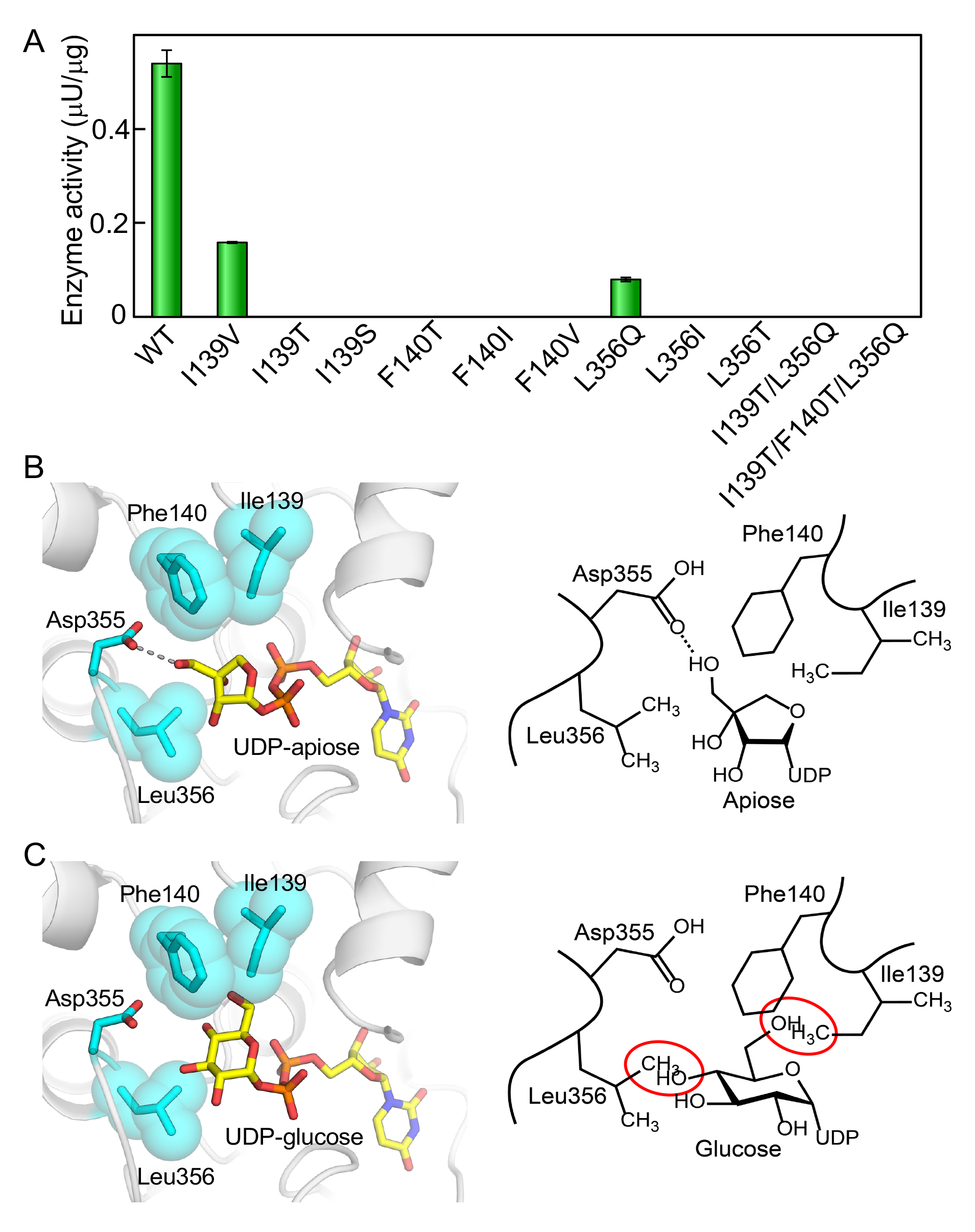
Ile139 and Leu356 are responsible for the specificity of AgApiT for UDP-Api. (A) Apiosyltransferase activities of wild-type (WT) and eleven AgApiT mutants are shown. The activities are presented as mean values with standard errors of three independent samples. (B) The structural model of AgApiT bound with UDP-Api (left). The sugar nucleotides and the side chains of some amino acid residues around the sugar portion are shown in stick form. The side chains of Ile139, Phe140, and Leu356 are also depicted as space-filling forms. The hydrogen bonds between Asp355 and 3′-OH of apiose residue are shown by dotted lines. Schematic representation of AgApiT recognizing UDP-Api (right). (C) Structural model of AgApiT tentatively bound with UDP-Glc (left) and its schematic representation (right). The space of 4-OH of glucose residue and the side chain of Leu356, or 6-OH of glucose residue and the side chain of Ile139 and Phe140 overlaps are shown by red circles. UDP-Glc is practically not recognized by AgApiT.

We further evaluated the role of Ile139 and Leu356 for enzyme activity by calculating enzyme kinetic parameters of I139V and L356Q mutants for sugar donor and acceptor substrates. Thus, for mutant I139V, the *K*_m_ and *k*_cat_ values for the sugar acceptor apigenin 7-*O*-glucoside were similar to those of the wild-type AgApiT (Table 3). Additionally, the *k*_cat_ value for sugar donor UDP-Api was similar to that of the wild-type AgApiT but the *K*_m_ value for UDP-Api was two times higher than that of wild-type AgApiT (Table 3). As for L356Q, the *K*_m_ value for UDP-Api was nearly 45-fold higher than that of the wild-type AgApiT. In turn, the *k*_cat_ value was also affected, being approximately 4.6-fold lower than that of the wild-type AgApiT (Table 3). Lastly, the *K*_m_ and *k*_cat_ values of mutant L356Q for the sugar acceptor substrate, apigenin 7-*O*-glucoside, were also affected, although not as much as for the sugar donor substrate. Altogether, these data indicated that at least Ile139 and Leu356 are involved in the recognition of UDP-Api by AgApiT.

**Table 3.** Kinetic parameters of wild-type, L356Q, and I139V of AgApiT.

| Substrate | Enzyme | $K_m$ ($\mu\text{M}$) | $k_{\text{cat}}$ ($\times 10^{-3} \text{ s}^{-1}$) | $k_{\text{cat}}/K_m$ ($\text{M}^{-1} \text{ s}^{-1}$) |
| --- | --- | --- | --- | --- |
| UDP-apiose | WT | $8.6 \pm 0.6$ | $0.65 \pm 0.01$ | $76 \pm 6$ |
| | I139V | $17 \pm 1$ | $0.47 \pm 0.01$ | $28 \pm 3$ |
| | L356Q | $390 \pm 60$ | $0.14 \pm 0.01$ | $0.36 \pm 0.08$ |
| Apigenin 7-*O*-glucoside | WT | $15 \pm 3$ | $0.88 \pm 0.05$ | $58 \pm 15$ |
| | I139V | $17 \pm 1$ | $0.83 \pm 0.07$ | $50 \pm 18$ |
| | L356Q | $52 \pm 9$ | $0.21 \pm 0.02$ | $2.1 \pm 0.3$ |

These values were calculated from the nonlinear curves obtained from three measurements at each concentration of individual substrates.

A previous study on VvGT1 revealed that the amino acid residue Thr141 forms a hydrogen bond with 6-OH of the glucose moiety of UDP-Glc in the crystal structure of VvGT1 (Offen et al., 2006). Similarly, the side chain of Ile139 was predicted to be located proximal to C-4 of the apiosyl residue in UDP-Api in the structural model of UDP-Api-bound AgApiT (Figure 5B). In addition, the side chain of Phe140 was also predicted to be located near C-4 or C-3 of the apiosyl residue. The side chain spaces of Ile139 and Phe140 likely occupy the site for the hydroxymethyl group at position 6 of the glucose moiety of UDP-Glc (Figure 5C), which might be one of the reasons why AgApiT recognizes UDP-Api but is inert to UDP-Glc.

In the structural model of AgApiT (Figure 5B), the terminal amino group of Asp355 is predicted to form hydrogen bonds with 3′-OH of apiosyl residue. In turn, the side chain of Leu356 is supposed to be close to C-2 and C-3 of apisoyl residue, such that the hydrophobic interaction between them may be a critical factor for the high specificity of AgApiT for UDP-Api. The space for the side chain of Leu356 likely occupies the site for the hydroxy group at C-4 of the glucose moiety of UDP-Glc or UDP-Xyl, suggesting a likely reason explaining why AgApiT is inert with UDP-Glc and UDP-Xyl (Figure 5C).

In the case of tea XylT (CsGT2/UGT94P1), the Ser substitution for Ile139 in AgApiT reportedly caused the loss of specificity for xylose and showed specificity for glucose instead (Ohgami et al., 2015). However, for AgApiT, neither GlcT nor XylT activity was detected in any of the nine amino acid‒substituted mutants at positions Ile139, Phe140, or Leu356, when UDP-Glc or UDP-Xyl was used as sugar donor substrates (Figure 5A). In the structural model of AgApiT, the space corresponding to 6-OH or 4-OH of the glucose moiety of UDP-Glc was filled with the side chain of either Ile139, Phe140 or Leu356 (Figure 5C). In addition, double (I139T/L356Q) and triple (I139T/F140T/L356Q) mutants substituted with the same amino acids most similar to GlcT (PgUGT94Q2) in amino acid sequence to AgApiT (Figure 1) lost ApiT activity (Figure 5A) and showed neither GlcT nor XylT activity. This spatial arrangement by side chains of at least three amino acid residues (Ile139, Phe140, and Leu356) in the substrate pocket of AgApiT would presumably contribute to the strict specificity of this enzyme for UDP-Api.

To further confirm the importance of these three amino acid residues for apiose recognition, we examined whether Agr35256-2 protein, the most homologous celery GGT to AgApiT with 60% sequence homology (Figure 1; Supplemental Figure S2), had ApiT activity. The Ile139, Phe140, and Leu356 residues in AgApiT were replaced by Val139, Asn140, and Gln361 in Agr35256-2, respectively (Supplemental Figure S2). We did not detect apiosyltransferase activity in the recombinant Agr35256-2 protein expressed in *E.coli* under the condition which AgApiT showed apiosyltransferase activity. This supports the idea that the Ile139, Phe140, and Leu356 residues in AgApiT are important for apiose recognition, and that AgApiT is the only apiin-synthetic apiosyltransferase in the celery genome.

## Discussion

AgApiT catalyzes the transfer of the apiosyl residue from UDP-Api to the 2-position of the Glc residue of flavone 7-*O*-glucosides to produce the apiosylated flavone including apiin (Figures 3 and 4). One of the key features of AgApiT is its strict sugar-donor specificity (Figure 4A) and relatively low *K*_m_ value for UDP-Api (Figure 4 and Table 3). By comparing amino acid sequences of GGTs that utilize UDP-Glc or UDP-Xyl as a specific sugar donor, we found unique amino acid residues near the UDP-Api binding site of AgApiT. Furthermore, site-directed mutagenesis studies based on homology modeling of AgApiT showed that Ile139, Phe140 and Leu356 were responsible for the specific recognition of the apiose residue of UDP-Api (Figure 5).

A molecular phylogenetic tree of GGTs from several plant species and 26 celery GGTs is shown in Figure 1. The GGTs with amino acid sequences most similar to AgApiT were Panax ginseng PgUGT94Q2, which is the GlcT for ginsenoside Rg_3_ and Rd (Jung et al., 2014), and sesame SiUGT94D1, which is the GlcT for the biosynthesis of sesaminol glucosides (Noguchi et al., 2008). Although they clustered in the same clade of the GGT phylogenetic tree (Figure 1), these two GlcTs show different acceptor and donor specificities from those of AgApiT, and do not contain Ile139, Phe140, or Leu356, which are crucial for apiose recognition by AgApiT. Moreover, among 26 celery GGTs, AgApiT is the only one possessing these three amino acid residues, suggesting that *AgApiT* is the only ApiT gene in the celery genome and that other highly-homologous paralogs are non-ApiTs. Sequential substitutions of several amino acid residues, including three amino acid residues identified in this study from the ancestral GGT enzyme, are likely a prerequisite for the evolution of ApiT. However, multiple substitutions at the unique three amino acid positions in AgApiT are not sufficient to revert the ancient sugar donor specificity of ApiT, suggesting that additional amino acid substitutions are involved in the evolution of ApiT. This notion is consistent with no UGT subclade with a specific sugar donor specificity (Figure 1) and sporadically distribution of glycosyltransferases with unique sugar donor specificities in the phylogeny of the UGT94 family. The UGT94 includes XylT (Ohgami et al., 2015), GlcT (Noguchi et al., 2008; Ono et al., 2010), glucuronyltransferase (Sawada et al., 2005), rhamnosyltransferase (Frydman et al., 2004), and ApiT (in this study).

AgApiT uses various flavonoid glucosides as sugar acceptor substrates. Indeed, it is highly predisposed to transfer apiose to the glucose attached to the 7-position of apigenin, quercetin, and luteolin, and less to those attached to naringenin (Figure 4B). Considering that the presence or absence of a hydroxyl group attached to the 3 or 3’ position of the flavone or flavonol skeleton is not relevant for recognition by AgApiT, the enzyme shows a high affinity for 7-*O*-glucosides with a flavone or flavonol skeleton and less specificity for those with a flavanone skeleton. Therefore, AgApiT should be more appropriately called flavone 7-*O*-glucoside: ApiT rather than apiin-biosynthesis ApiT. This name has been defined already in EC 2.4.2.25 as the enzyme name UDP-Api:7-*O*-(β-D-glucosyl)-flavone apiosyltransferase (Ortmann et al., 1972). This sugar acceptor preference of AgApiT is consistent with the distribution of apiose-containing flavonoid glycosides in celery. Apigenin (54%), chrysoeriol (23%), and luteolin (23%) glycosides have been detected in celery, and more than 99% of them are apiosylated (Lin et al., 2007). Thus, we speculate that this single enzyme catalyzes the biosynthesis of all these apiosylated compounds in celery by exerting catalytic promiscuity for sugar acceptors.

In contrast to sugar acceptor specificity, AgApiT has a relatively low *k*_cat_ value and high affinity for UDP-Api. (Table 3). This may result from the acquisition of apiose specificity at the expense of metabolic turnover efficiency under the neofunctionalization of the enzyme. The three-dimensional structural analysis of AgApiT will increase our understanding of enzymatic evolvability and provide structural insights into substrate specificity and catalytic efficiency of GGT for sugar donors and acceptors in the future.

Apiin was initially found in the Apiaceae (including celery and parsley) as a specialized flavonoid glycoside. It is sporadically observed in phylogenetically distant families, including the Asteraceae, Fabaceae, and Solanaceae families, suggesting convergent evolution of specificity to UDP-Api from different UGT genes. Thus, it is challenging to identify ApiT genes from other plant families based on total amino acid sequence similarity in the context of AgApiT. Ile139, Phe140, and Leu356 residues that recognize apiose may provide clues in identifying them.

It should be noted that AgApiT (GT1 family) identified in this study is unrelated to bacterial ApiT, which belongs to the GT90 family (Smith and Bar-Peled, 2018). The acceptor substrate of this bacterial ApiT remains to be clarified. Comparing the spatial arrangement of the amino acid‒side chains for apiose recognition of plant and bacterial ApiT warrants further exploration.

The celery *CHS, CHI,* and *FNSI,* encoding the enzymes for apigenin biosynthesis are co-expressed in the early leaf growth stages (Yan et al., 2014) (Supplemental Figure S1). Their regulatory transcription factor genes have also been reported (Yan et al., 2019; Wanget al., 2022). Gene expression analysis of *AgApiT* during various developmental stages of celery showed that its expression pattern is similar to that of *FNSI* (Figure 2). This spatio-temporal coordination suggests that *AgApiT* is also co-regulated with these apiin-biosynthetic enzymes by common transcription factors, thereby orchestrating apiin biosynthesis during the early leaf-developmental stages in celery. Therefore, our results confirm that spatio-temporal co-expression is a reliable footprint to reveal the biosynthetic genes in the same specialized metabolic pathway (Chae et al., 2014).

Our findings provide a biotechnological platform for heterologously producing apiin in systems such as *E. coli*. In this case, co-expression of AgApiT with UDP-Api biosynthetic enzymes such as AXS (Mølhøj et al., 2003) may be effective. In turn, the efficient production of apiin may help understand the biological activity and the benefits of apiin or apiose residues for human health in the future. Finally, the identification of AgApiT provides an opportunity to generate apiin-deficient or apiin-accumulating plants to assess the physio-ecological roles of apiin in nature.

## Materials and Methods

### Plant materials

The celery (cultivar ‘New Cornell 619’) seeds were purchased from Takii Seed (Kyoto, Japan). The seeds were grown on MS medium at 22°C under a 16-hour light/8-hour dark cycle. After 10 days, the seedlings were transferred to a soil mixture of Metro-Mix 360 (Sun Gro Horticulture, Agawam, OH) and vermiculite (4:1) under the same conditions.

### GGT genes from celery

Celery RNA-Seq dataset SRR1023730 was obtained from the Sequence Read Archive (SRA) of NCBI (https://www.ncbi.nlm.nih.gov/sra). The reads were trimmed using Trimmomatic and *de novo* assembled using Trinity on the Galaxy server (http://usegalaxy.org). The reads were mapped using HISAT2, and the Fragments per Kilobase Megareads (FPKM) values were calculated for each predicted gene using StringTie. Celery GGTs (UGT79, UGT91, and UGT94) sequences were obtained using BLAST among the transcripts and predicted genes in the celery genome database (Li et al., 2020) using known UGT sequences as queries.

### Phylogenetic analysis

The sequences of plant GGTs (UGT79, UGT91, and UGT94) biochemically characterized so far were obtained from the UGT Nomenclature Committee Website (https://labs.wsu.edu/ugt/). The sequences of celery GGTs were obtained as described above. These sequences were assembled with ClustalW, and their phylogenetic tree was constructed using the neighbor-joining method.

### Heterologous expression of AgApiT

The AgApiT and Agr35256-2 ORFs containing restriction enzyme sites were synthesized chemically as codon-optimized genes (Eurofins Scientific, Luxembourg) for protein expression in *E. coli.* The genes were amplified with [AgApiTpCold_F and R] and [Agr35256-2_F and R] primer sets (Supplemental Table S1) and cloned into the NdeI/XbaI sites of the pCold ProS2 vector (Takara Bio, Kusatsu, Japan). The resulting plasmid was transformed into *E. coli* BL21 (DE3). The cells were cultured at 37°C till they reached an OD_600_ of 0.6, after which, 0.8 mM isopropyl β-D-thiogalactoside was used to induce the expression of recombinant AgApiT fused with the proS2 tag for 24 h at 15°C. Then, cells were harvested by centrifugation at 5,000 × *g* and 4°C for 5 min and lysed with BugBuster Protein Extraction Reagent (Merck Millipore, Burlington, MA) prepared in 20 mM sodium phosphate buffer (pH 7.4) and supplemented with 5 U/mL benzonase and 1 kU/mL lysozyme. Cell debris was removed by centrifuging the lysate at 20,000 × *g* for 10 min at 4°C. The resulting supernatant was loaded onto a 1 mL-HisTrap HP column (Cytiva, Marlborough, MA) equilibrated with buffer A (50 mM sodium phosphate buffer and 500 mM sodium chloride, pH 7.4) containing 50 mM imidazole. The column was washed with buffer A containing 100 mM imidazole, and the protein was eluted with the same buffer containing 200 mM imidazole. The fusion protein was digested with HRV3C protease at 4°C for 2 h to remove the proS2 tag. The resulting AgApiT protein was purified as a flow-through fraction on a 1 mL-HisTrap HP column equilibrated with buffer A containing 50 mM imidazole. The protein was concentrated by ultrafiltration using an Amicon Ultra-0.5 mL (10 kDa cutoff, Merck Millipore) at 14,000 × *g* and 4°C for 15 min. Protein concentration was determined using a Pierce 660 nm protein assay kit (Thermo Fisher Scientific, Waltham, MA) and protein quality was analyzed by SDS-PAGE followed by Coomassie Blue staining.

### Site-directed mutagenesis of AgApiT

*In vitro* mutagenesis was performed by PCR using specific mutagenesis oligonucleotide primers (Supplemental Table S1) and the KOD-Plus Neo (Toyobo, Osaka, Japan) with pColdProS2-AgApiT (6,288 bp) as the template. The following site-directed mutants were obtained: I139S, I139T, I139V, F140I, F140T, F140V, L356I, L356Q, and L356T. The plasmid template in the resulting PCR product was digested using DpnI, and each of the mutants was transformed into *E. coli* DH5α cells. Individual clones of these *E. coli* variants were grown, and corresponding plasmids were isolated. Then, individual mutations were verified by DNA sequencing. All mutant proteins were expressed and purified using the same procedures as those used for the wild-type AgApiT. The double (I139T/L356Q) and triple (I139T/F140T/L356Q) mutant proteins were also prepared in the same manner as described above.

### Glycosyltransferase assay

Apiin and apigenin 7-*O*-β-D-glucoside were purchased from Ark Pharm (Arlington Heights, IL). Apigenin was purchased from Cayman Chemical (Ann Arbor, MI). Naringenin 7-*O*-β-D-glucoside, luteolin 7-*O*-β-D-glucoside, and quercetin 7-*O*-β-D-glucoside were obtained from Extrasynthese (Genay, France). Chrysoeriol 7-*O*-β-D-glucoside was obtained from ChemFaces (Wuhan, China). UDP-Api (Fujimori et al., 2019), UDP-Xyl (Ishimizu et al., 2005), UDP-rhamnose (Rha) (Ohashi et al., 2016), and UDP-galacturonic acid (GalUA) (Ohashi et al. 2006) were prepared as described. UDP-arabinofuranose (Ara*f*) was purchased from Peptide Institute (Osaka, Japan). UDP-Glc, UDP-galactose (Gal) and UDP-*N*-acetylglucosamine (GlcNAc) were purchased from Fuji Film Wako Chemical Corporation (Osaka, Japan). UDP-glucuronic acid (GlcUA) and GDP-fucose (Fuc) were purchased from Sigma-Aldrich. The glycosyltransferase assay was conducted using 50 μM acceptor substrate, 1 mM sugar nucleotide, and the purified recombinant protein in 200 mM Tris-HCl buffer containing 50 mM NaCl (pH 7.0) at 23°C for 2 h. The reaction was stopped by incubation at 100°C for 3 min. The substrate and the product were separated by reversed-phase HPLC using an Inertsil ODS-3 column (4.6 × 250 mm, GL Sciences, Tokyo, Japan) at a flow rate of 1.0 mL/min with an isocratic flow of 20% acetonitrile containing 0.1% trifluoroacetic acid for initial 5 min, followed by a linear gradient from 20 to 40% acetonitrile for 20 min. Apigenin, chrysoeriol, luteolin, naringenin, and quercetin were detected and quantified based on absorbance at 330, 330, 350, 280, and 254 nm, respectively. One unit of enzyme activity was defined as the amount of the enzyme that produced 1 µmol of the product per minute under the reaction conditions described above. The *K*_m_ and *k*_cat_ values for AgApiT were determined by assaying various concentrations of apigenin 7-*O*-β-D-glucoside (5–150 μM) or UDP-Api (2.5–150 μM). The kinetic parameters were calculated from the Michaelis-Menten equation by nonlinear regression analysis using Prism version 9. (GraphPad Software, San Diego, CA). The experiments were performed in triplicate to obtain the mean, and S.D. was shown as error bars.

### Molecular modeling of AgApiT

The initial homology model of AgApiT was constructed with MODELLER 10.2 (Šali et al. 1993) using the crystal structure of UGT71G1 bound with UDP-Glc (2acw) (Shao et al., 2005), VvGT1 (2c1x) (Offen et al., 2006), UGT85H2 (2pq6) (Li et al., 2007), UGT72B1 (2vce) (Brazier-Hicks et al., 2007), and UGT78G1 (3hbf) (Modolo et al., 2009) as templates. To generate the UDP-Api complex with AgApiT, the homology model of AgApiT was superimposed on UGT71G1 bound with UDP-Glc, and then the sugar portion of UDP-Glc was manually replaced with apiose. Energy minimization and structure equilibration were performed for the AgApiT complex bound with UDP-Api under the CHARMM force field (Brooks et al., 2009) using GROMACS 2020.1 (Berendsen et al., 1995).

### RT-PCR

The developmental leaves of celery were divided into seven stages according to the methodology proposed by Yan et al. (2014); stage 1: true leaves <0.5 cm in length; stage 2: between 0.5 and 1.0 cm; stage 3: between 1.0 and 1.5 cm; stage 4: between 1.5 and 2.0 cm; stage 5: between 2.0 and 2.5 cm; stage 6: between 2.5 and 3.0 cm; stage 7: >3.0 cm. These plant materials were immediately frozen in liquid nitrogen and stored at −80°C until use. Semi-quantitative reverse transcription PCR was performed in each developmental stage as follows. RNA was extracted using RNeasy plus mini kit (Qiagen, Hilden, Germany) and then reverse transcribed into cDNA using PrimeScript II 1^st^ strand cDNA synthesis kit (Takara Bio). PCRs for *AgApiT* and *FNS I* (Yan et al. 2014) were performed with a cDNA template using specific primer sets (Supplemental Table S1). Glyceraldehyde-3-phosphate dehydrogenase (GAPDH) (Gao and Loescher, 2000) was used as an internal control to verify that equal amounts of RNA were used from each sample. The quantity of PCR products was maintained within the linear range.

### Accession Number

The DNA sequence of *AgApiT* (*UGT94AX1*) has been deposited in DDBJ/ENA/GenBank under the accession number LC704571.

### Funding

This work was supported by a Grant-in-Aid for Scientific Research (18H05495, 19H03252, and 20K21403) from the Ministry of Education, Culture, Sports, Science and Technology of Japan. It was also supported by the Fugaku Foundation and the Program for the Fourth-Phase R-GIRO Research from the Ritsumeikan Global Innovation Research Organization, Ritsumeikan University.

